# Principled clustering of functional signals reveals gradients in processing both within the anterior hippocampus and across its long axis

**DOI:** 10.1101/2022.02.08.479587

**Authors:** John N. Thorp, Camille Gasser, Esther Blessing, Lila Davachi

## Abstract

A particularly elusive puzzle concerning the hippocampus is how the structural differences along its long, anteroposterior axis might beget meaningful functional differences, particularly in terms of the granularity of information processing. One measure posits to quantify this granularity by calculating the average statistical independence of the BOLD signal across neighboring voxels, or inter-voxel similarity (IVS), and has shown the anterior hippocampus to process coarser-grained information than that in the posterior hippocampus. This model of the hippocampus, however, conflicts with a number of task-oriented findings, many of which have varied in their fMRI acquisition parameters and hippocampal parcellation methods. In order to reconcile these findings, we measured IVS across two separate resting-state fMRI acquisitions and compared the results across many of the most widely used parcellation methods in a large young-adult sample (Acquisition 1, N = 253; Acquisition 2, N = 183). Finding conflicting results across acquisitions and parcellations, we reasoned that a principled, data-driven approach to hippocampal parcellation is necessary. To this end, we implemented a group masked independent components analysis (mICA) to identify functional subunits of the hippocampus, most notably separating the anterior hippocampus into separate anterior-medial, anterior-lateral, and posteroanterior-lateral components. Measuring IVS across these components revealed a decrease in IVS along the medial-lateral axis of the anterior hippocampus but an increase from anterior to posterior. We conclude that representational granularity may not change linearly or unidirectionally across the hippocampus, and that moving the study of the hippocampus towards reproducibility requires grounding it in a functionally informed approach.

**Significance Statement:** Processing information along hierarchical scales of granularity is critical for many of the feats of cognition considered most human. Recently, the changes in structure, cortical connectivity, and apparent functional properties across parcels of the hippocampal long axis have been hypothesized to underlie this hierarchical gradient in information processing. We show here, however, that the choice of parcellation method itself drastically affects the perceived granularity across the hippocampus, and that a principled, functionally informed approach to parcellation reveals gradients both within the anterior hippocampus and in non-linear form across the long axis. These results point to the issue of parcellation as a critical one in the study of the hippocampus and reorient interpretation of existing results.

---

It is often taken to be a key challenge for our memory system to extract and store general knowledge of the world while simultaneously retaining details of the individual episodes that make it up. For example, after seeing a number of shows at the jazz club downtown, one is able to call upon generalized representations to predict what the next set will be like and identify similar music on the radio, all while being able to precisely reconstruct where one sat during a particular performance on a rainy night two years ago. Recent work has proposed that the processing of information at these hierarchical levels of detail may be dependent on distinct functional specializations along the long axis of the hippocampus (Poppenk, Evensmoen, Moscovitch, & Nadel, 2013; Sekeres, Winocur, & Moscovitch, 2018; Strange, Witter, Lein, & Moser, 2014). Specifically, it has been argued that hippocampal representations become increasingly finer grained along the long axis, with integrative, coarse-grained representations in the anterior end and differentiated, fine-grained representations in the posterior end.

As a direct test of this proposed gradient, Brunec & Bellana et al., (2018) measured the average temporal correlation across voxels, termed inter-voxel similarity (IVS), as a putative index of the ‘granularity’ of hippocampal information processing. Mathematically, IVS is the mean correlation between BOLD activity timecourses of all voxels within a given region of interest (Fig 1A). Lower IVS values therefore indicate that the temporal activation profiles in voxels within a particular anatomical region are relatively uncoupled from each other, which has been taken to reflect more distinct information processing in adjacent voxels and therefore greater representational granularity. Using this measure, Brunec & Bellana et al., (2018) reported that, during virtual navigation as well as during rest, the anterior hippocampus displayed higher IVS than the posterior hippocampus, suggesting that the anterior hippocampus may support representations that are intrinsically coarser-grained than in its posterior counterpart.

**Figure 1.**
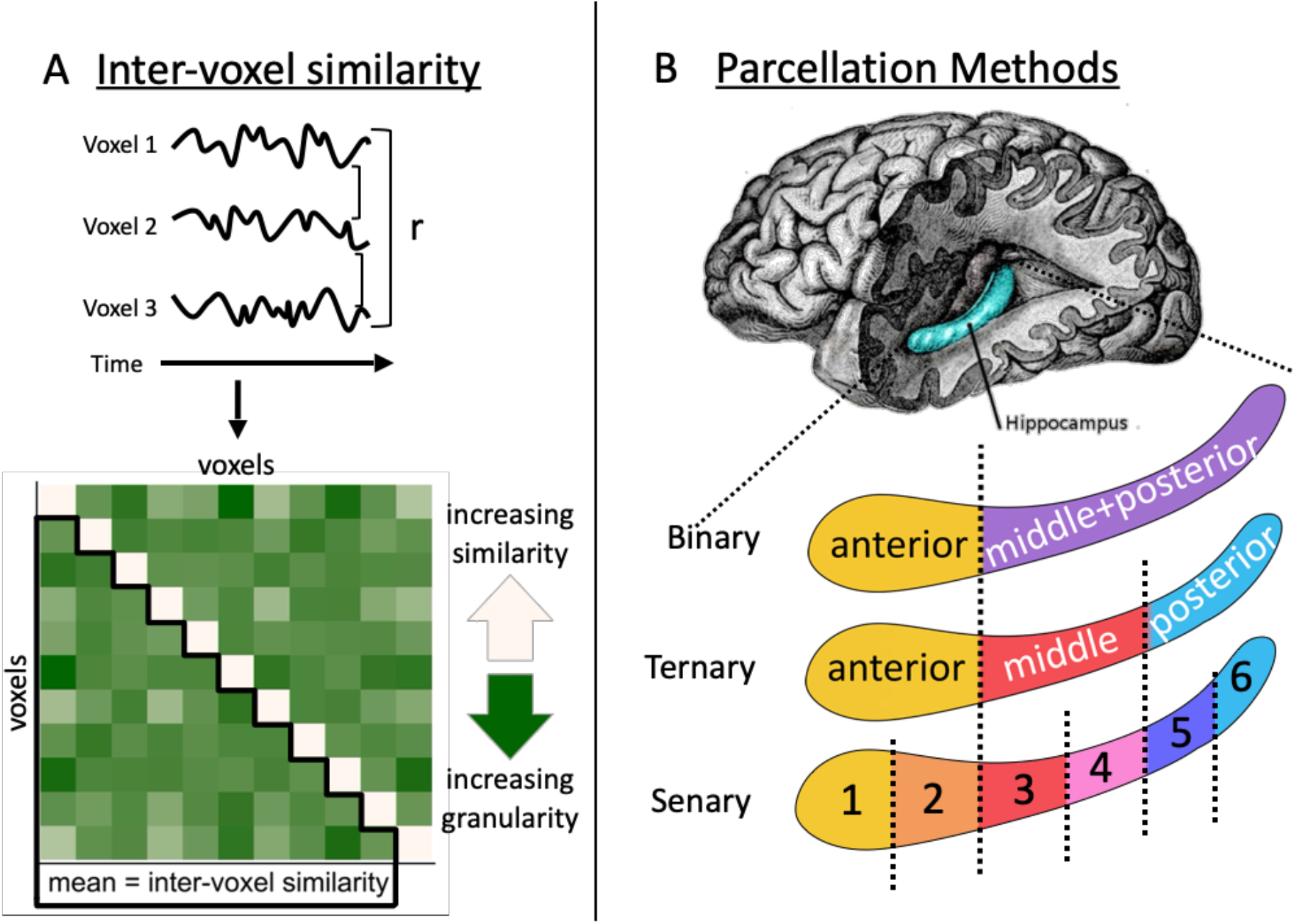
Overview of Methods. (**A**) Diagram of inter-voxel similarity (IVS), the mean pairwise correlation of every voxel’s entire timeseries within a given parcel of the hippocampus. Decreases in IVS are taken as indications of individual voxels being sensitive to more distinct information, and therefore processing representations with a higher granularity. (**B**) Proportional parcellations split the hippocampus (top) at roughly the uncal apex (binary parcellation), into even thirds (ternary parcellation), or into even sixths (senary parcellation).

Interestingly, however, other applications of IVS along the long axis have found inconsistent results in both developmental (Callaghan et al., 2020) and healthy aging (Stark, Frithsen, & Stark, 2021) samples: namely, that IVS in these populations increases from anterior to posterior hippocampus. These findings that patterns of hippocampal granularity may not be static across the lifespan are complemented by additional work indicating those patterns may differ across task demands as well (Brunec, Robin, Olsen, Moscovitch, & Barense, 2020). For example, representational similarity analyses comparing evoked BOLD patterns across items have shown that the anterior hippocampus may differentiate representations of individual items (Ezzyat, Inhoff, & Davachi, 2018; Ranganath & Ritchey, 2012) while posterior hippocampus has been shown to integrate representations of items paired with the same scene (Tompary & Davachi, 2017). These conflicting results raise the question as to whether IVS is a stable measure of any essential properties of information processing.

A major roadblock in reconciling these findings, however, is the variation in parcellation methods used to divide the hippocampus into its constituent subregions. Researchers have most typically divided the hippocampal long axis into binary, ternary, and senary (Fig 1B) parcellation methods, the former two of which fall in line with structural and genetic transitions, respectively (Poppenk et al., 2013; Strange et al., 2014). While much of the literature has assumed the choice of parcellation method to be relatively trivial, thorough understanding of its impact is critical to proper interpretation of functional measures like IVS.

In the current study, we first used a large resting-state sample across two fMRI acquisition sequences (Acquisition 1, N = 253; Acquisition 2, N = 183) to compare the pattern of IVS across the hippocampus when using canonical parcellation methods (i.e. binary, ternary, senary). While we should expect these proportional parcels to provide a rough summary of the differences across the long axis, it is unlikely to be the case that they have perfectly localized the natural joints of the emergent functional subunits we aim to study. Therefore, we adopted an additional analytical approach in which we applied group masked independent components analysis (mICA) to the resting-state signals in the hippocampus, which essentially identifies the spatial map that maximizes the statistical independence across individual components (Blessing, Beissner, Schumann, Brünner, & Bär, 2016; Blessing et al., 2020; Moher Alsady, Blessing, & Beissner, 2016). We then compared IVS across these functionally-derived parcels, stepping closer to a true characterization of functional gradients across the hippocampus.

## Materials and Methods

### fMRI Dataset

Analyses were performed on the Nathan Kline Institute-Rockland Sample (NKI-RS), a lifespan cross-sectional dataset obtained at the Nathan Kline Institute and made publicly available online (Nooner et al., 2012). Participants were scanned on a SIEMENS 3T MRI scanner. A high-resolution 3D MPRAGE T1-weighted anatomical (TR = 1900 ms, voxel size = 1 mm isotropic, FoV = 250 mm) was first obtained, followed by two separate resting state scans with differing acquisition parameters. The first resting state scan (TR = 2500 ms, TE = 30 ms, voxel size = 3 mm x 3 mm x 3.5 mm, duration = 5 minutes, FOV = 216 mm) is referred to as Acquisiton 1, and the second resting state scan (TR = 1400 ms, TE = 30 ms, voxel size = 2 mm isotropic, duration = 10 minutes, FOV = 216 mm, multi-band accel. factor 4) referred to as Acquisition 2 (Table 1). Participants were only instructed to keep their eyes open and fixate on the screen for the duration of each scan.

**Table 1.** Scan Acquisition Parameters across Studies

| Acquisition | TR<br>(ms) | TE<br>(ms) | voxel size<br>(mm) | duration<br>(min) | FOV |
| --- | --- | --- | --- | --- | --- |
| Brunec & Bellana et al., (2018) | 2000 | 24 | 3.5 x 3.5 x 3.5 | 6 | 20 cm<sup>2</sup> |
| Acquisition 1 | 2500 | 30 | 3 x 3 x 3.5 | 5 | 216 mm |
| Acquisition 2 | 1400 | 30 | 2 x 2 x 2 | 10 | 216 mm |

*Note:* Acquisition 1 more closely aligns with the scan parameters employed during the resting-
state analysis in Brunec & Bellana et al., (2018).

### fMRI Preprocessing

The fMRI preprocessing steps we applied are in line with the pipeline used by Brunec & Bellana et al., (2018), with the exception that Statistical Parametric Mapping 12 (SPM12) was used rather than SPM8. In Acquisition 1, functional images were slice-time corrected, realigned and resliced in accordance with the mean functional image, and coregistered to the space of the anatomical images. Acquisition 2 was preprocessed identically except that slice-time correction was not performed due to its faster TR during multi-band acquisition. The anatomical images were then segmented into cerebro-spinal fluid (CSF) and white matter (WM) images, which were thresholded at 0.7 and 0.9 (Menon & Uddin, n.d.), respectively, and eroded. These masks were used to extract CSF and WM timeseries from the functional images. The six motion regressors extracted from realignment as well as the CSF and WM timeseries were then entered as multiple regressors in a denoising generalized linear model (GLM).

### Conservative motion and BOLD interpolation

The residuals of this GLM (the denoised functional timeseries) were then fed through a data interpolation method based on principal components analysis (PCA) (Brunec et al., 2018; Campbell, Grigg, Saverino, Churchill, & Grady, 2013). Because our analysis of the heterogeneity of individual voxel timeseries is sensitive to overly liberal interpolation, the method employed here only interpolates a timepoint if both its motion regressors and BOLD signal are significantly far from their own medians (Campbell et al., 2013). First, PCAs were run on both the matrix of time x six motion regressors and the matrix of time x voxels. Then, for each timepoint, the squared, normalized distance between the timepoint and the median of the surrounding 15 timepoints was calculated, resulting in a vector of distances from the median for each timepoint for each matrix. A gamma distribution was then estimated for each vector, and the cumulative density function estimated for each timepoint. Timepoints that had a p < 0.05 in both the six motion regressors and the functional timeseries were interpolated using spline interpolation. In other words, only timepoints that were significant outliers in both their overall motion and BOLD signal were removed and interpolated using surrounding data points.

### Proportional parcellations

To define a mask for each participant’s hippocampus, anatomical images were segmented into subcortical structures using the recon-all function in FreeSurfer (version = 7.1.1) (Fischl, 2012). These subcortical atlases were then transformed back into native space and the bilateral hippocampus masks were extracted.

In order to mirror hippocampal parcellation approaches adopted by previous studies, we divided the hippocampus into subregions along the long axis by following three separate parcellation methods. For our binary parcellation, the hippocampus was divided into an anterior third plus a combined middle+posterior two-thirds (Fig 1B, left). This parcellation serves to approximate that used by Brunec & Bellana et al., (2018), who split the individual hippocampi at the uncal apex into anterior and posterior parcels, and is similar to previous specifications (Brunec et al., 2020; Poppenk et al., 2013). For our ternary parcellation, we divided the hippocampus into even anterior, middle, and posterior thirds along the long axis (Callaghan et al., 2020; Collin, Milivojevic, & Doeller, 2015; Dandolo & Schwabe, 2018; Tompary & Davachi, 2017). Finally, we also split the hippocampus into even sixths in a senary parcellation (Stark et al., 2021).

### Masked Group Independent Components Analysis

While the proportional parcellations described above might offer a rough approximation of unidimensional changes in granularity across the long axis, they also rest on top-down assumptions about how to identify meaningful hippocampal subunits. As such, also sought to parcellate the hippocampus based on the observed structure of activity within its constituent voxels (i.e., in a data-driven. Scans from the 183 participants (for Acquisition 2) were submitted to a mICA using FSL Melodic implemented with the mICA toolbox (version=1.18). In the masked group ICA, a dimensionality of 10 was chosen based on previous research showing 10 components displays the highest split-half reproducibility without over-parcellating the hippocampus (Blessing et al., 2016).

### Extended fMRI preprocessing for group mICA

Due to multi band slicing artifacts present in Acquisition 2, data were first cleaned using the fix FSL package (version=1.06) (Griffanti et al., 2014; Salimi-Khorshidi et al., 2014). The fix FSL package essentially functions by isolating individual components intrinsic to the data using Melodic ICA, classifying components as noise or signal, and regressing noise components out of the data. We specifically implemented the training dataset WhII_MB6 that comes with the fix FSL package that closely aligned with the scan parameters from Acquisition 2, and used the default threshold of 20% confidence to binarize signal from noise. These cleaned functional images were only used for creating the group parcellation, and IVS was still calculated using the original images.

The functional images cleaned using the fix FSL package were then smoothed using a smoothing kernel with a full-width half-maximum of 6 mm. A high pass filter was then applied to the functional images with a cut-off of 100-secs. Functional images from all participants were then normalized to the MNI avg152 T1-weighted template (2 mm isotropic resolution).

### Group masked ICA Parcellation

We then implemented the group masked ICA, which concatenated all 183 functional images (from Acquisition 2) and masked them with the HarvardOxford bilateral hippocampal mask thresholded at 50%, as in previous studies (Blessing et al., 2016, 2020; Moher Alsady et al., 2016). Melodic ICA, with a dimensionality of 10, was then performed as previously described. Briefly, the data were demeaned and normalized by the voxel-wise variance, projected into a 10-dimensional space using probabilistic PCA, and decomposed into spatial maps and timeseries using a fixed-point iteration technique optimizing for non-Gaussian spatial distribution. The resulting group-level spatial maps of each component were then divided by the standard deviation of the residual noise and thresholded with a mixture model fitted to the histogram of each component to yield a z-transformed spatial map for each component. This process therefore yields the spatial maps of 10 functionally derived parcels of the hippocampus.

### Warping group masked ICA parcels into native space

In order to use these spatial maps as individual masks for parcellating the hippocampus, each component was manually labeled based on the five locations within the hippocampus, thresholded at 0.5, and binarized. As the components from the ICA are not necessarily spatially cohesive (i.e., they are not constrained to one contiguous region), all voxels that were not spatially contiguous with the area containing the maximum z-score were erased using FreeView’s edit function. The component masks were then warped back into native space. To better align the components with the hippocampus in native space, the full Harvard-Oxford hippocampus mask was also warped into each participant’s native space and linearly coregistered with the FreeSurfer hippocampal mask used before. The registration matrix retrieved from this step was then applied to each of the components, ensuring that each component was as closely aligned to its true location in native space as possible.

In order to confirm that the components derived from the mICA were indeed aligned properly within the FreeSurfer hippocampal mask, we calculated the proportion of the component mask that overlapped with the FreeSurfer mask for each participant and component. Any participant’s component not overlapping at least 20% with the FreeSurfer mask was excluded from further analyses, though results were qualitatively similar without this exclusion.

### Extracting parcel timeseries

Voxel-wise timeseries were then extracted from the functional image using the hippocampal masks. This procedure produced a single timeseries for each voxel, scan, and participant in each of our hippocampal parcels.

### Controlling for parcel size and shape

To control for the variable sizes and shapes of the parcels, the average distances between the location of each voxel and the center voxel was calculated separately for each of the x, y, and z axes within each parcel, acquisition, and participant, as performed in Brunec & Bellana et al., (2018). These average distances were then converted into units of millimeters along each dimension and later entered as parcel-level covariates in subsequent mixed effects models.

### Inter-voxel similarity calculation and exclusion criteria

All timeseries analyses were conducted using R (version = 3.5.0) and RStudio. A correlation matrix for each participant, hemisphere, and hippocampal parcel was constructed by first z-scoring the timeseries within each voxel, and then finding the correlation of each voxel’s timeseries with every other voxel’s timeseries within a given mask. Using original code from Brunec & Bellana et al., (2018), the resulting correlation matrices were then Fisher-z transformed, the diagonal and upper triangle removed, and the mean of the resulting vector computed. This procedure resulted in one value of IVS per hippocampal parcel, acquisition, and participant. In order to protect against signal drop-out, any hippocampal parcel missing data from more than 30% of voxels was also excluded. Only scans with greater than 0.56 mm mean frame displacement were excluded. Finally, individual outlier values of IVS were excluded by first grouping values by acquisition, hemisphere, and parcel. Values that were farther than 1.5 times the inter-quartile range of this distribution from its median were assumed to be due to scanner noise and excluded from further analyses. This left an N of 235 participants (118 females, 1 undisclosed) in Acquisition 1, and an N of 183 participants (85 females) in Acquisition 2.

## Statistical Analyses

### Mixed Effects Models Predicting Inter-Voxel Similarity

Linear mixed effects models were performed using the lmer() function from the lme4 package (Bates, Mächler, Bolker, & Walker, 2015) in R. Separate models were run per acquisition. For models used to compare ternary and binary parcellation methods, IVS was entered as the dependent variable; hemisphere (effect-coded “left” = −0.5, “right” = 0.5), parcel (“anterior”, “middle”, “middle+posterior”, “posterior”, reference = “anterior”), and their interactions were entered as independent variables. Models accounting for parcel size and shape included x_mean_distance, y_mean_distance, and z_mean_distance as covariates (see “Controlling for parcel size and shape” for detail). All models included a random intercept grouped by participant, as was done in Brunec & Bellana et al., (2018).

For our senary parcellation of even sixths, one model was used to extract the parcel-wise standard error estimates shown in Figure 4, while another was used to find the linear slope presented in Figure 4. Note that the parcel-wise standard error estimates are only presented for visual purposes, and the only statistical analyses shown are the linear slopes. The first model was identical to those described above, except for the inclusion of six rather than four levels within the parcel variable (“one”, “two”, “three”, “four”, “five”, “six”, reference = “one”). Mean distances along the x, y, and z axes were included as covariates. The model used to calculate the slope of the line of best fit along these parcels was the same as the one above except that the parcel factors were converted to their quantitative equivalents (1, 2, 3, 4, 5, 6, respectively) and mean-centered. Again, mean distances along the x, y, and z axes were included as covariates. In order to estimate p values for the calculated slopes, degrees of freedom were approximated for each fixed effect using the same Kenward-Roger method as for the estimated marginal means above.

The final mixed effects model was used to examine how IVS changed between the components derived from the mICA, and was the same as the others above except for the factors within the parcel variable (“anterior-medial”,”anterior-lateral”, “posteroanterior-lateral”, “middle”, posterior”, reference = “anterior-medial”). Again, mean distance along the x, y, and z axes were entered as covariates.

In order to directly compare IVS values across hippocampal parcels within hemisphere, the emmeans() function (Lenth, 2020) in R was used to extract estimated marginal means for each parcel averaged across hemisphere from each model as well as run pairwise t-tests between parcels. Degrees of freedom were approximated using the Kenward-Roger method (Kenward & Roger, 1997), and p-values were adjusted via Tukey’s method for comparing a family of estimates of the given size (Tukey, 1949).

## Code Accessibility

Code for analyzing IVS and recreating figures will be accessible on Github.

## Results

Our first goal was to investigate how inter-voxel similarity (IVS) changes along the long axis of the hippocampus, while specifically considering how these results differ across the multiple hippocampal parcellation methods often adopted by researchers in this field. We began by comparing IVS across binary (anterior vs. middle+posterior) and ternary (anterior vs. middle vs. posterior) parcellations, performing linear mixed effects regressions with a random intercept per participant, mirroring the primary analysis from Brunec & Bellana et al., (2018). In order to directly compare IVS across hippocampal parcels, pairwise t-tests were run between estimated marginal means of IVS computed for each parcel averaged across hemisphere. The Tukey method was used to correct for multiple comparisons within a family of four estimates (anterior, middle, middle+posterior, and posterior). Separate mixed effects models were run for Acquisition 1 and Acquisition. Note that Acquisition 1 more closely aligns with the scan parameters implemented in Brunec & Bellana et al., (2018), while Acquisition 2 provides many more voxels, TRs, and overall timepoints within each participant (Table 1).

### Inter-voxel similarity decreases from anterior to posterior hippocampus

We first aimed to replicate Brunec & Bellana et al., (2018) by directly comparing IVS within the anterior parcel (yellow) to that in the middle+posterior parcel (purple) across acquisitions (Fig 2A). We found that IVS within the anterior parcel was significantly higher than that in the middle+posterior parcel during both Acquisition 1 (t(1442) = 15.00, p_tukey_ < 0.001) and Acquisition 2 (t(1156) = 18.95, p_tukey_ < 0.001), which indeed mirrors the pattern observed during both resting-state and spatial navigation in Brunec & Bellana et al., (2018).

**Figure 2.**
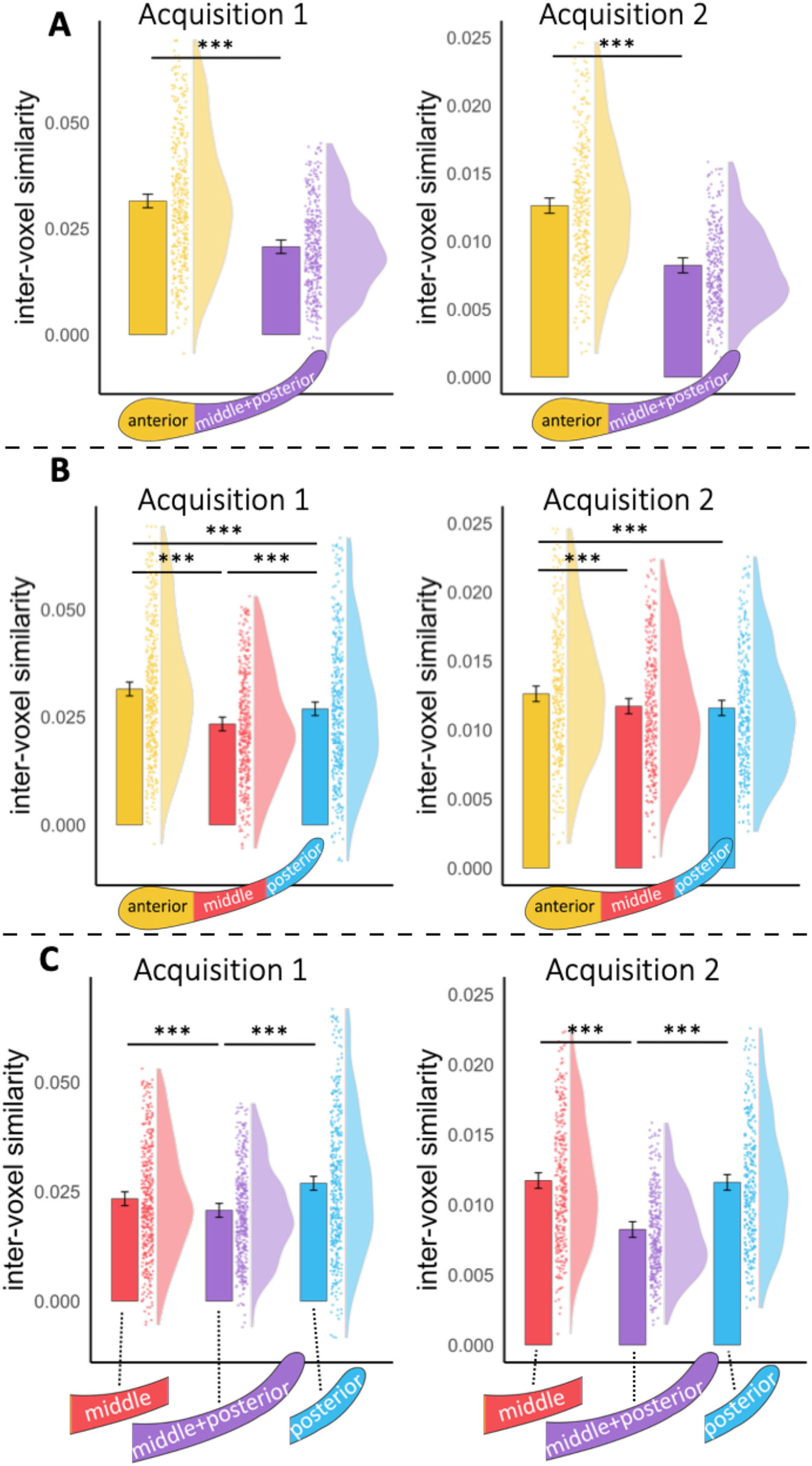
Binary parcellation underestimates the inter-voxel similarity of the middle and posterior subregions. (**A**) We replicate the key result of Brunec and Bellana (2018) that inter-voxel similarity (IVS) within the anterior parcel (yellow) is higher than that in the middle + posterior parcel (purple) across both acquisitions. (**B**) Upon splitting the middle+posterior parcel into its constituent middle (red) and posterior (blue) parcels, as has been done elsewhere, IVS seemed to be higher within the anterior parcel than that in the posterior parcel across both acquisitions, though interestingly IVS within the middle parcel was lower than that in both the anterior and posterior parcels in Acquisition 1, in conflict with a linear decrease along the long axis. (**C**) Critically, however, the middle+posterior parcel displayed a significantly lower IVS than both its constituent middle and posterior regions across both acquisitions, suggesting that combining these subregions into one misrepresents their apparent granularity. This last finding motivated our following analyses, which take the overall parcel size and shape into account.

Next, we computed IVS within ternary parcels (e.g. (Callaghan et al., 2020; Collin et al., 2015; Dandolo & Schwabe, 2018) by splitting the middle+posterior parcel into its constituent parts (middle [red] and posterior [blue] subregions). As would be predicted by the prior result, IVS within the anterior third parcel remained higher than that in the posterior third parcel during Acquisition 1 (t(1445) = 6.40, p_tukey_ < 0.001) as well as Acquisition 2 (t(1156) = 4.45, p_tukey_ < 0.001) (Fig 2B).

### U-shaped change in inter-voxel similarity along ternary parcels

Prior work examining the hippocampus with this ternary approach has most typically provided evidence that the middle hippocampus appears as a linear combination of what is seen in the anterior and posterior hippocampus (Callaghan et al., 2020; Collin et al., 2015; Dandolo & Schwabe, 2018). Thus, one strong prediction might be that IVS should smoothly decrease from anterior to middle to posterior hippocampus. Following this prediction, we found that IVS within the anterior parcel was significantly higher than that in the middle parcel in during Acquisition 1 (t(1440) = 11.33, p_tukey_ < 0.001) as well as Acquisition 2 (t(1153) = 3.90, p_tukey_ < 0.001) (Fig 2B). However, in contrast to what would has been predicted by a linear gradient in granularity, IVS within the middle parcel was also lower than that in the posterior parcel during Acquisition 1 (t(1433) = −5.01, p_tukey_ < 0.001), though not during Acquisition 2 (t(1152) = 0.58, p_tukey_ = 0.94).

### The middle+posterior parcel exaggerates the granularity of its constituent subregions

To further unravel the systematic differences between these binary and ternary parcels, we next considered in the relationship between the apparent granularity measured within the combined middle+posterior parcel and that of its constituent middle and posterior parcels (when calculated separately). We found that the middle+posterior parcel consistently displayed lower IVS than that in the middle parcel during both Acquisition 1 (t(1428) = −3.78, p_tukey_ < 0.001) and Acquistion 2 (t(1151) = −15.34, p_tukey_ < 0.001) as well as the posterior parcel during both Acquisition 1 (t(1429) = −8.80, p_tukey_ < 0.001) and Acquisition 2 (t(1150) = −14.76, p_tukey_ < 0.001) (Fig 1C).

This pattern of results suggests that combining these potentially distinct subregions into the same parcel and correlating their voxel timeseries together may artificially lower the apparent IVS within the unified parcel. In other words, it may be that the low IVS within the middle+posterior region is more directly caused by the mixing of disparate signals from middle and posterior hippocampus rather than the individual voxels in this subregion being relatively uncoupled from each other. Not only does this overaccentuate the difference in granularity between the anterior and posterior subregions, it also glosses over what may be a more complex U-shaped gradient in processing from anterior to middle to posterior hippocampus.

### Accounting for parcel size and shape accentuates U-shaped change in ternary parcels

One way to account for the middle+posterior parcel’s overestimation of its own granularity is to add control variables for the overall size and shape of each parcel. Since we would expect the timeseries of voxels anatomically farther away from one another to be less similar to each other, a parcel that spans a longer distance along the hippocampus may display a lower IVS simply by nature of including voxels anatomically farther apart. Thus, we implemented a control analysis from Brunec & Bellana et al. (2018) that simply adds covariates to each model to account for the spatial size of each parcel along the x, y, and z axes. Unstandardized betas for each covariate are reported in Table 2. The resulting model estimates therefore more precisely represent the decoupling of individual voxel timeseries from one another, controlling for the average anatomical distance between them.

**Table 2.**
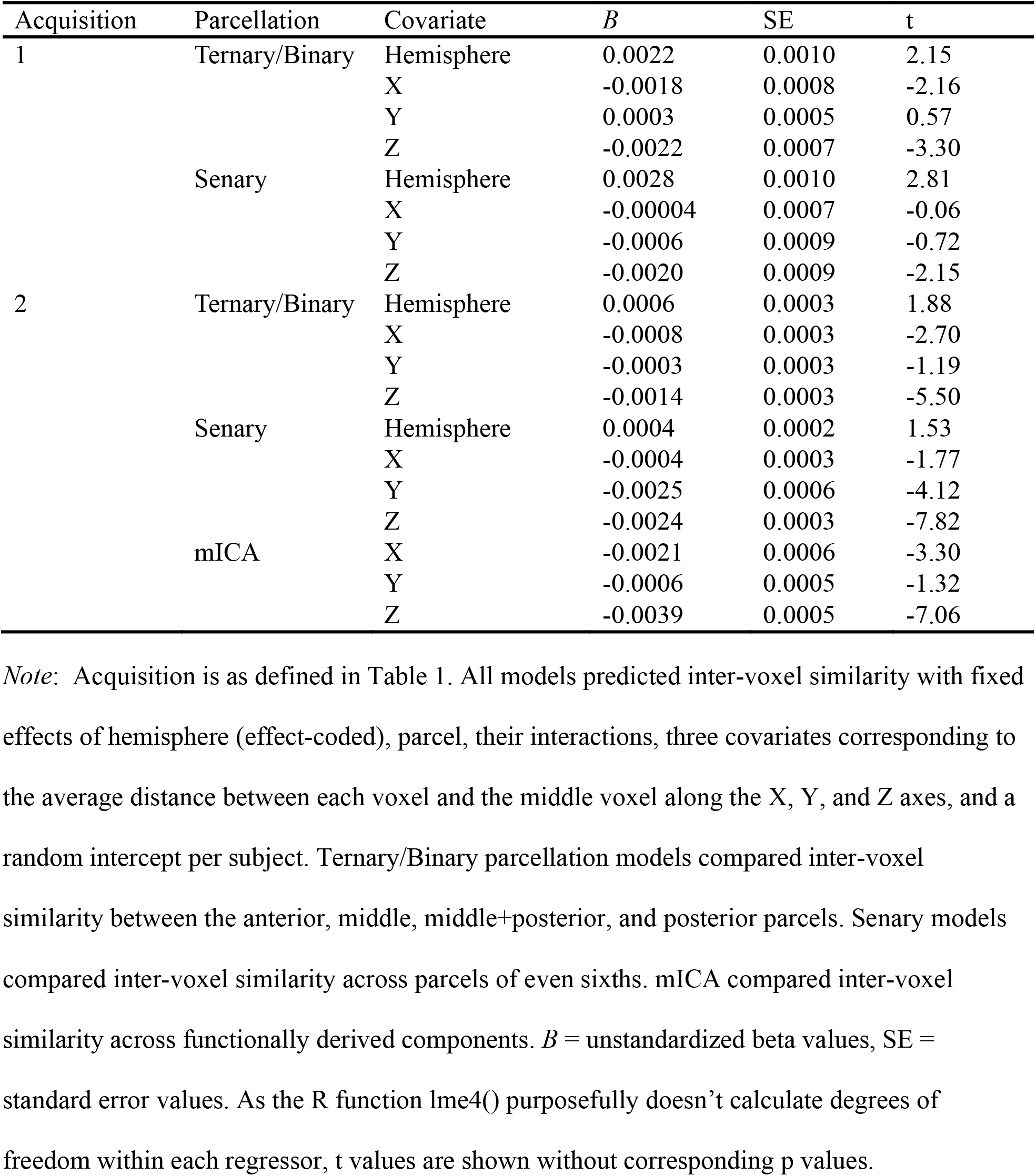
Influence of model covariates on inter-voxel similarity

While IVS in the anterior parcel remained higher than that in the middle+posterior parcel during Acquisition 1 (t(1535) = 4.16, p_tukey_ < .001), consistent with Brunec & Bellana et al., (2018), this difference was not statistically significant during Acquisition 2 (t(1249) = 1.05, p_tukey_ = .72) (Fig 3A). Similarly within our ternary parcels, while IVS in the anterior parcel remained higher than that in the posterior parcel during Acquisition 1 (t(1562) = 3.78, p_tukey_ < .001), this difference was again not statistically significant during Acquisition 2 (t(1273) = 1.44, p_tukey_ = .47) (Fig 3B). Across both parcellation methods, therefore, apparent differences in granularity between anterior and posterior hippocampus are left intact during Acquisition 1 but well accounted for by parcel size and shape during Acquisition 2.

**Figure 3.**
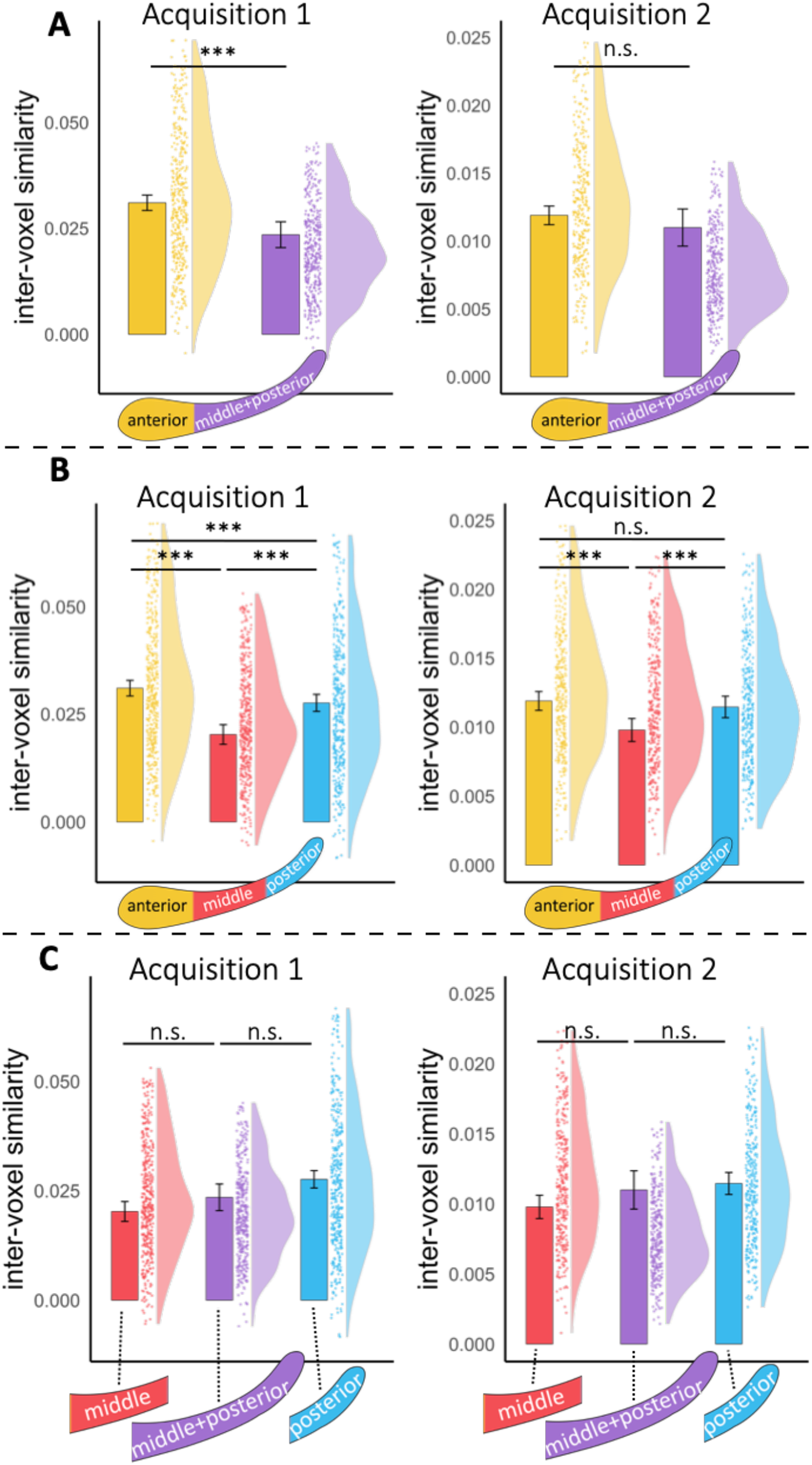
Controlling for parcel size and shape reinforces a U-shaped change in granularity along the long axis. To account for the discrepancies in length, width, and height between the different parcels, covariates for parcel size and shape were added to the models, as was accounted for in a later analysis in Brunec & Bellana et al., (2018). (**A**) After entering these covariates, we still found substantial evidence for the original comparison between anterior and middle+posterior in Acquisition 1, but not in Acquisition 2. (**B**) In our ternary parcels, the anterior parcel displayed a higher IVS than that in the posterior parcel in Acquisition 1, but not in Acquisition 2. Across both acquisitions, however, IVS in the middle parcel was lower than that in both the anterior and posterior parcels, reinforcing the U-shaped change along the long axis. (**C**) As was the intended result of adding these covariates, no differences were found between the middle+posterior parcel and its constituent middle and posterior parcels, as these were completely accounted for by their differences in size and shape.

In contrast, the addition of these parcel size and shape covariates only served to strengthen the U-shaped change in IVS from anterior to middle to posterior hippocampus. That is, IVS in the middle parcel remained lower than that in the anterior parcel in both Acquisition 1 (t(1579) = −10.14, p_tukey_ < .001) and Acquisition 2 (t(1282) = −5.81, p_tukey_ < .001), as well as lower than that in the posterior parcel in both Acquisition 1 (t(1608) = −6.00, p_tukey_ < .001) and Acquisition 2 (t(1305) = −3.75, p_tukey_ = .001). As anticipated, the addition of these covariates also accounted for the differences between middle+posterior and its constituent middle and posterior parcels (all p_tukey_’s > 0.14) (Fig 3C). Taken together, these results across acquisitions seem to suggest that accounting for parcel size and shape reinforces a U-shaped gradient in IVS along the hippocampal long axis.

### Inter-voxel similarity increases along senary parcels of even sixths

To follow up on these results at a finer resolution, we performed mixed effects linear regressions examining IVS across senary parcels of even sixths. As above, this model included fixed effect covariates of mean distance along the x, y, and z axes and a random intercept grouped by subject. Directly conflicting with the findings in Stark et al., (2021) in young adults, we found evidence of a linear increase along the long axis (from anterior to posterior) in both Acquisition 1 (*B* = 0.0013, t(2205) = 4.00, p_tukey_ < .001) as well as Acquisition 2 (*B* = 0.00033, t(1799) = 3.80, p_tukey_ < .001). In other words, modeling IVS linearly using this senary parcellation shows that IVS is lowest (more granular) in anterior hippocampus and is highest in posterior hippocampus. These inconsistent results across parcellation methods and acquisitions should suggest to us that slicing unilaterally across the hippocampus may not be carving at the most relevant joints. That is, just as the middle+posterior parcel may combine signals across disparate middle and posterior subregions, it may be the case that any of our typical, proportional parcellation methods inadvertently combine data across disparate clusters of functional activity.

### Group masked independent components analysis

Our results thus far demonstrate that the proportional parcellation techniques largely adopted by the field fall short of characterizing the true functional subunits of the hippocampus. As such, we next aimed to parcellate the hippocampus based on the underlying structure of the functional signals themselves. To this end, we ran a group masked independent components analysis (mICA) (Blessing et al., 2016, 2020; Moher Alsady et al., 2016) that effectively separates the hippocampus into the 10 components that maximize the spatial independence between components and the temporal coherence within components. Due to the comparatively higher number of TRs in Acquisition 2, only Acquisition 2 was submitted to the mICA and used in the following set of analyses, which benefit greatly from more within-subject data. This procedure resulted in the bilateral hippocampi being clustered into ten distinct regions, five in each hemisphere: three splitting the anterior hippocampus (anterior-medial [AM, red], anterior-lateral [AL, yellow], and posteroanterior-lateral [PAL, green]), one in the middle hippocampus (middle [M, light blue]), and one in the posterior hippocampus (posterior [P, dark blue]) (Fig 5A). Split-half reproducibility analyses resulted in an average Pearson’s correlation of 0.966 across components, showing that randomly splitting the sample into discrete halves had virtually no effect on the spatial layout of the components.

**Figure 4.**
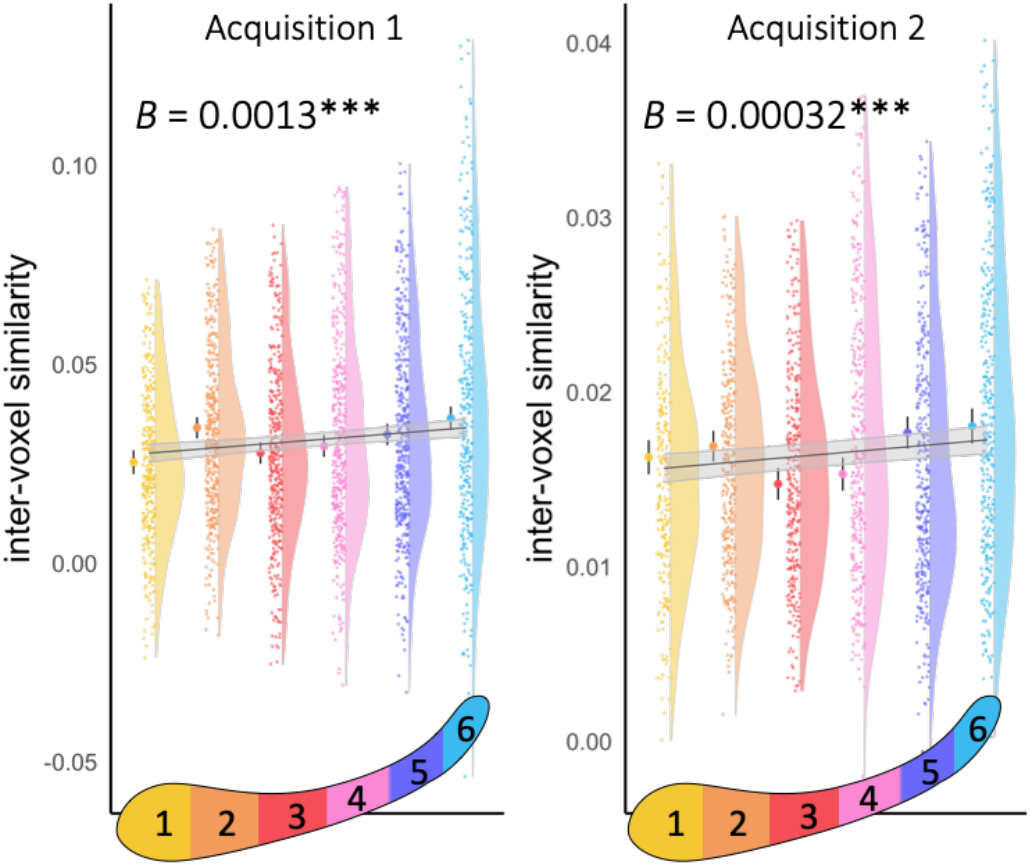
Inter-voxel similarity increases across parcels of even sixths. When modeled linearly, inter-voxel similarity shows a significant increase from anterior to posterior across parcels of even sixths in both acquisitions.

**Figure 5.**
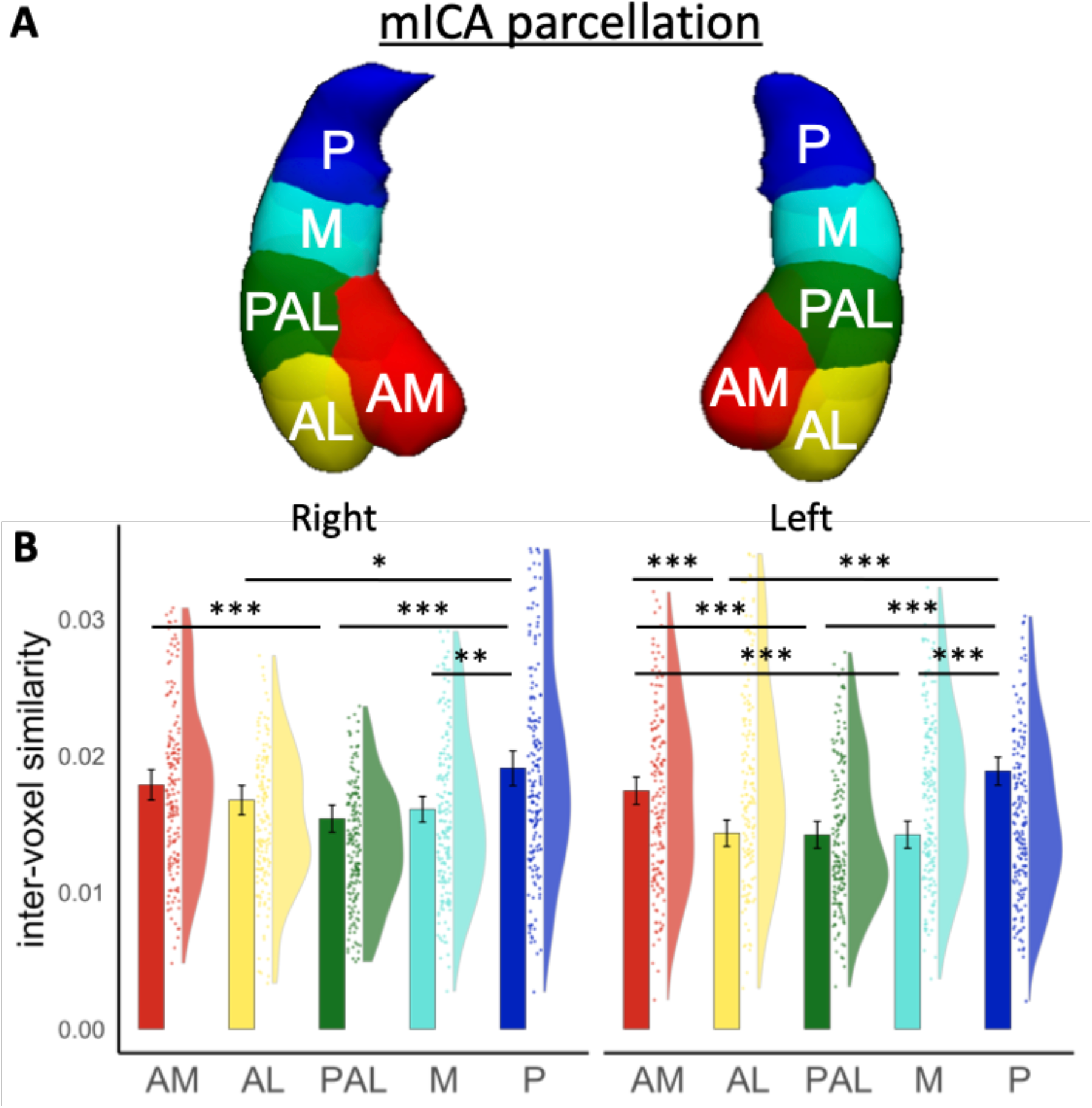
Principled parcellation of functional signals reveals a medial-lateral gradient of inter-voxel similarity within the anterior hippocampus. (**A**) Parcellation as derived from group masked ICA. Replicating earlier work, the mICA split each hemisphere into anterior-medial (AM, red), anterior-lateral (AL, yellow), posteroanterior-lateral (PAL, green), middle, (M, light blue), and posterior (P, dark blue) components. (**B**) Because of differences in parcellation across hemispheres, we report differences in inter-voxel similarity in both hemispheres. Contrary to our findings using proportional parcellations, inter-voxel similarity was higher within the posterior parcel than that in the anterior-lateral, posteroanterior-lateral, and middle parcels across both hemispheres. Interestingly, inter-voxel similarity within the anterior-medial parcel was higher than that in the middle, posteroanterior-lateral, and anterior-lateral parcel within the left hemisphere, and higher than that in the posteroanterior-lateral parcel in the right hemisphere. This suggests that processing granularity may differ along the medial-lateral axis within the anterior hippocampus, a difference that was inaccessible using the proportional parcellations used previously.

In order to gain a better understanding of how granularity, as measured by IVS, might differ across these functional subregions, we next calculated IVS within each component and ran a linear mixed-effects model predicting IVS by hippocampal component and hemisphere. As above, this model included fixed effect covariates of mean distance along the x, y, and z axes and a random intercept grouped by subject. Due to the fact that the mICA creates slightly different parcels across hemispheres, we performed all t-tests within hemisphere, implementing the Tukey method to correct for a family of 5 tests in each hemisphere.

### mICA finds a medial-lateral gradient within anterior hippocampus

When looking at these functionally derived components, we again found evidence of a U-shaped gradient in IVS between the anterior-medial component (red) and the posterior component (blue) (Fig 5B). IVS within the posterior parcel was higher than that in the anterior-lateral component (yellow) in both the left (t(1536) = 7.62, p_tukey_ < 0.001) and right (t(1603) = 2.94, p_tukey_ = 0.028) hemispheres, the posteroanterior-lateral component (green) in both the left (t(1568) = 7.36, p_tukey_ < 0.001) and right (t(1566) = 5.31, p_tukey_ < 0.001) hemispheres, as well as the middle component (light blue) in both the left (t(1550) = 7.66, p_tukey_ < 0.001) and right (t(1595) = 3.80, p_tukey_ = 0.001) hemispheres. IVS within the anterior-medial component (red) was then higher than that in the middle component in the left (t(1504) = 5.57, p_tukey_ < 0.001) and right (t(1567) = 2.89, p_tukey_ = 0.032) hemispheres.

Critically, we also found a decrease in IVS along the medial-lateral axis of the anterior hippocampus. That is, IVS within the anterior-medial component was higher than that in the posteroanterior-lateral component in both the left (t(1519) = 4.96, p_tukey_ < 0.001) and right (t(1513) = 4.20, p_tukey_ < 0.001) hemispheres, as well as than that in the anterior-lateral component in the left hemisphere (t(1492) = 5.28, p_tukey_ < 0.001), though this difference was not significant in the right hemisphere (t(1619) = 1.42, p_tukey_ = 0.61). These differences suggest what could be an important axis within the anterior hippocampus that has been left unexamined by typical proportional parcellation methods.

## Discussion

Inter-voxel similarity (IVS) is a measure of the statistical similarity of voxels within a given subregion. The intuition behind this measure is that subregions responsible for extracting features that overlap across episodes, objects, or larger temporal windows would contain voxels that were responsive to similar information and therefore more temporally intertwined with one another. The degree of this temporal similarity has thus been used to suggest that intrinsic dynamics of information processing across the long axis of the hippocampus move from coarse-grained (high IVS) in the anterior to fine-grained (low IVS) in the posterior. Here, we used a large fMRI resting-state dataset to show that change in IVS along the long axis varies widely with the adopted parcellation method and seems to change along multiple gradients within the hippocampus.

We find that IVS decreases along binary hippocampal parcels from anterior to middle+posterior (Fig 7, ‘Binary’), in line with prior published reports. However, we also found that considering the middle and posterior thirds of the hippocampus as one ‘middle+posterior’ region artificially lowers the apparent IVS, ultimately suggesting that the IVS within this binary parcellation is not a reliable estimate of the true underlying signal variance. Accounting for the size and shape of the individual parcels then resulted in a U-shaped change in IVS along ternary parcels, with the middle parcel displaying lower IVS than that in the anterior and posterior parcels (Fig 7, ‘Ternary’). A senary parcellation into even sixths further complicated the story, with IVS displaying a linear increase from anterior to posterior hippocampus (Fig 7, ‘Senary’), thereby suggesting the posterior hippocampus is actually coarser-grained than its anterior counterparts – as has been found in both developmental (Callaghan et al., 2020) and healthy aging (Stark et al., 2021) cohorts.

**Figure 7.**
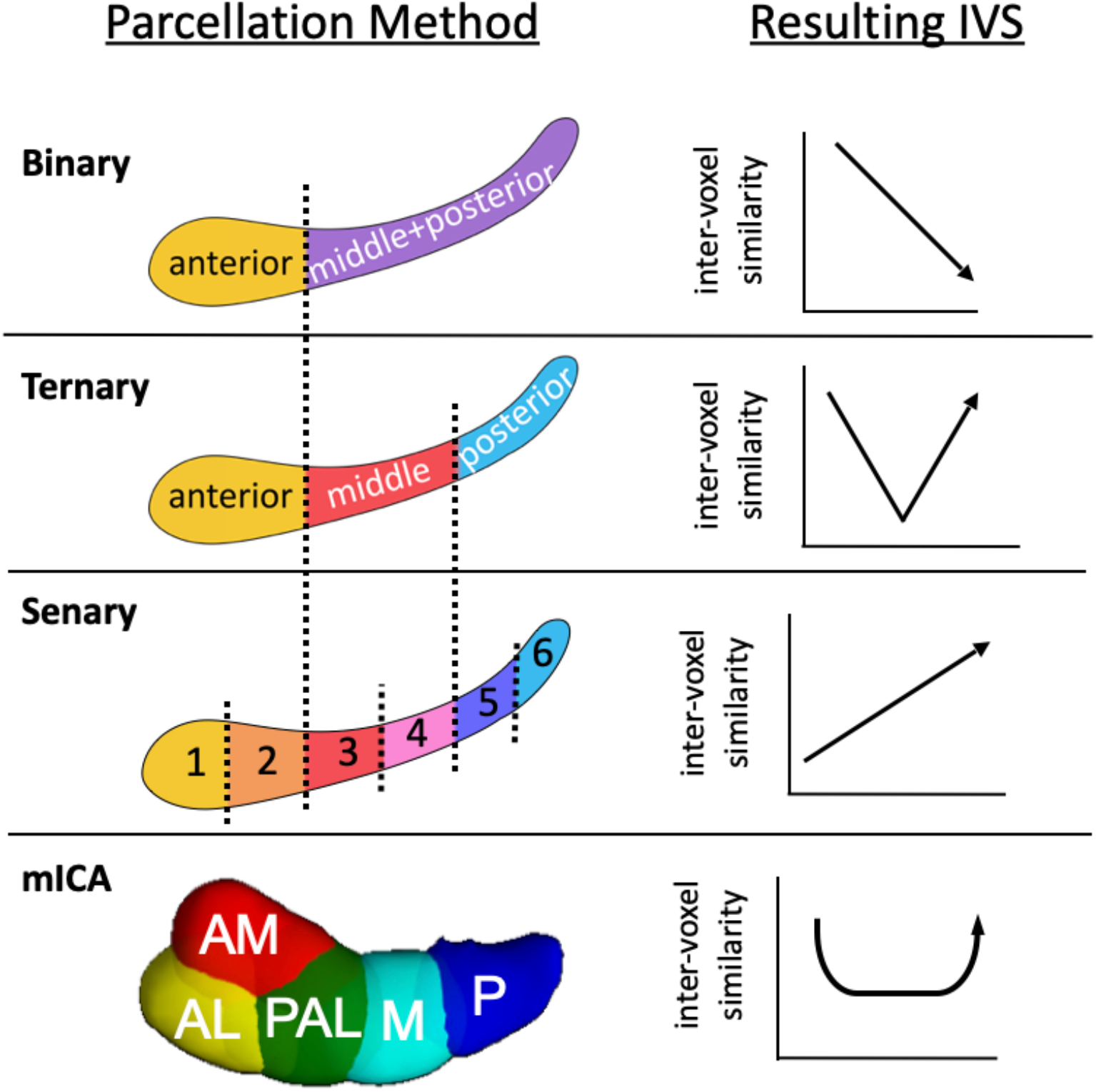
Summary of inter-voxel similarity findings within all parcellation methods. Inter-voxel similarity decreased along binary parcels, formed a U-shaped gradient along ternary parcels, increased along senary parcels of even sixths, and within the mICA parcellation both decreased along the medial-lateral axis within the anterior hippocampus and formed a U-shaped gradient from anterior-medial to posterior.

Thus, in order to take a step back, we came to the conclusion that parcellations drawn onto structural landmarks (i.e. the binary parcellation at the uncal apex) or genetic boundaries within the rodent hippocampus (i.e. the ternary parcellation) were not localizing the functional joints of the hippocampus. In order to more carefully localize these subunits before attempting to characterize them, we employed a group masked independent components analysis (mICA) to identify hippocampal subregions that contain voxels with similar temporal coherence. This data-driven parcellation revealed subregions analogous to those found across prior published studies (Blessing et al., 2016, 2020; Moher Alsady et al., 2016). Although this technique by definition will identify hippocampal clusters with relatively high levels of IVS, it is agnostic to differences in temporal coherence across distinct components – our measure of interest. Critically, this approach revealed a novel differentiation within the anterior hippocampus along its medial-lateral axis as well as U-shaped change from anterior to posterior, with IVS within the anterior-lateral and posteroanterior-lateral components lower than that in both the anterior-medial and posterior components (Fig 7, ‘mICA’).

The principal takeaway from our findings, then, is simply that hippocampal IVS is deeply affected by parcellation. More generally, that carving the hippocampus in a manner unrelated to the underlying functional clustering fails to accurately represent its functional characteristics. Though the majority of the literature has assumed the issue of parcellation to be relatively inconsequential, here we provide evidence it may be clouding our insights into true functional differences (Bryce et al., 2021). Thus, it seems best to ground one’s approach in a framework that doesn’t rely on genetic or structural signposts, but rather the emergent organization of the functional signals themselves. Such approaches of identifying meaningful boundaries between regions of the brain have proved extremely successful in the realm of cortical network segmentation (Beckmann, DeLuca, Devlin, & Smith, 2005; Smitha et al., 2017), but have yet to be the default method in intra-regional parcellation. Of the concerted effort to parcellate the hippocampus in a principled manner (Barnett, Man, & McAndrews, 2019; Chase et al., 2015; Cheng, Zhu, Zheng, Liu, & He, 2020; Robinson et al., 2015; Voets et al., 2014; Wang, Ritchey, Libby, & Ranganath, 2016; Zarei et al., 2013), most of the resulting methods have relied on clustering second-order consequences of the voxel timeseries (e.g. the IVS matrices used here, hippocampo-cortical connectivity, univariate results across studies) and have often replicated the proportional parcellations already typical to the literature.

The mICA parcellation is therefore somewhat unique in dealing directly with the voxel activity timeseries themselves, repeatedly uncovering a medial-lateral distinction within the anterior hippocampus that seems to rest on the localization of the uncus within the anterior-medial component (Zeidman & Maguire, 2016). Crucially, the formation of the uncus – via the rostromedial inversion of the anterior hippocampus during embryonic development – is distinctive to primates, such that the strict genetic demarcations found in the ventral hippocampus in rodents may not be representative of any such demarcations in the primate hippocampus (Strange et al., 2014). This medial-lateral distinction resonates with divergent findings where the anterior hippocampus may, at times, integrate overlapping item pairs during online concept formation (Bowman & Zeithamova, 2018; Davis, Love, & Preston, 2012a, 2012b; Mack, Love, & Preston, 2018; Viganò & Piazza, 2021) or immediate memory retrieval (Libby, Reagh, Bouffard, Ragland, & Ranganath, 2018; Ritchey, Montchal, Yonelinas, & Ranganath, 2015), while, at other times, differentiate between specific items in memory (Cowan et al., 2021; Ezzyat et al., 2018; Tompary & Davachi, 2017). It may be that these distinct medial-lateral components help to resolve this conflict.

It’s also worth noting the relatively consistent appearance of a U-shaped change in IVS from anterior to posterior, both across ternary parcels as well as our mICA components. While we resist interpreting any this too deeply without evidence of behavioral implications or validation of IVS across other measures, they raise an interesting question of whether it’s appropriate to consider the middle hippocampus as simply a linear midpoint between the anterior and posterior hippocampus on our measures of interest (Callaghan et al., 2020; Collin et al., 2015; Dandolo & Schwabe, 2018). Future work is needed to explicitly probe the role of this functional unit of the hippocampus within the distributed memory system.

## Limitations

As with all resting-state analyses, our results cannot fully explain how hippocampal activity maps onto behavior or information processing generally. Measuring brain activity during sophisticated behavioral tasks design is critical to the full characterization of the myriad roles of the hippocampus in the human brain. Rich indices of behavior and other neural measures of integration would also allow for a true appraisal of what computations or representations IVS might be measuring, which might reinform existing results.

Here we adopt the logic that low IVS may indicate more granular information processing, such that would underly coding of specific contextual or visuospatial details in memory representations. The strongest interpretations of IVS extend this measure to reflect the essential, intrinsic dynamics of the human hippocampus, ostensibly invariant to behavioral task demands or representational content. We resist that level of interpretation here. That is, we do not predict that the patterns of IVS shown above would hold across every behavioral task, type of attended content, or motivational state the hippocampus might represent or be involved in. To us, the evidence here simply shows how sensitive neural measures are to one’s prior organizational assumptions, and how complex and heterarchical the hippocampus appears when those assumptions are abandoned.

## Acknowledgements

This work was supported by National Institutes of Health Grant R01-MH-076492 to L.D. We thank Anna MacKay-Brandt and Michael Milham for assistance in accessing the imaging data, as well as Catalina Yang for refining a color palette suitable for the study of the long axis.

